# A closer look at cross-validation for assessing the accuracy of gene regulatory networks and models

**DOI:** 10.1101/190157

**Authors:** Shayan Tabe-Bordbar, Amin Emad, Sihai Dave Zhao, Saurabh Sinha

## Abstract

Cross-validation (CV) is a technique to assess the generalizability of a model to unseen data. This technique relies on assumptions that may not be satisfied when studying genomics datasets. For example, random CV (RCV) assumes that a randomly selected set of samples, the test set, well represents unseen data. This assumption does not hold true where samples are obtained from different experimental conditions, and the goal is to learn regulatory relationships among the genes that generalize beyond the observed conditions. In this study, we investigated how the CV procedure affects the assessment of methods used to learn gene regulatory networks. We compared the performance of a regression-based method for gene expression prediction, estimated using RCV with that estimated using a clustering-based CV (CCV) procedure. Our analysis illustrates that RCV can produce over-optimistic estimates of generalizability of the model compared to CCV. Next, we defined the ‘distinctness’ of a test set from a training set and showed that this measure is predictive of the performance of the regression method. Finally, we introduced a simulated annealing method to construct partitions with gradually increasing distinctness and showed that performance of different gene expression prediction methods can be better evaluated using this method.

## INTRODUCTION

A common method for computational reconstruction of gene regulatory networks (GRNs) is to build ‘expression-to-expression’ models, which predict gene expression as a function of expression levels of other genes, or of transcription factors (TFs) in particular (Fig. 1a); regulatory relationships are then learned from the parameters of such models ^1‒7^. This approach typically creates an over-parameterized model-fitting problem, since any subset of the hundreds to thousands of TFs coded in the genome may be regulators of a gene, and without prior knowledge of these regulators, the relationship between every TF and putative target gene must be encoded as a free parameter in the model. Different methods adopt various strategies to handle the resulting ‘curse of dimensionality’, and since a ‘ground truth’ GRN is almost never available to allow for an unbiased assessment of these methods, studies usually apply K-fold cross-validation (CV) for evaluation ^2–6^. This involves dividing the samples – experimental conditions under which expression of a gene (and all TFs) is known – into *K* disjoint subsets or ‘folds’, training the model’s free parameters on *K-1* of these folds, using the model to predict gene expression in the left-out samples, and finally comparing the predictions with measured values (Fig. 1b). This is repeated *K* times, so that all data samples (conditions) are used as a test sample once. Finally, by averaging the accuracy of predictions across all *K* repetitions, an overall measure of performance is obtained.

**Figure 1.**
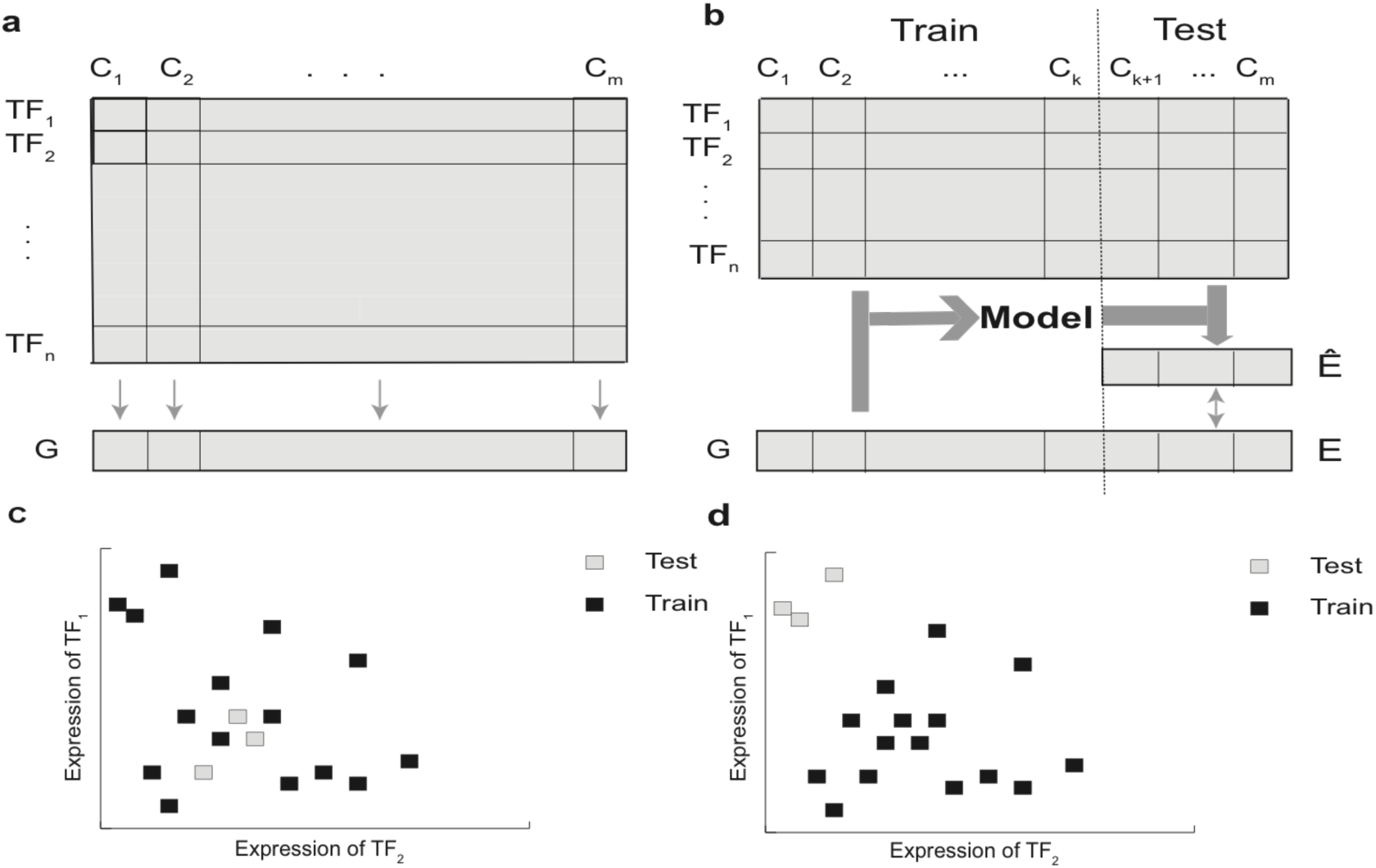
Schematic illustration of expression-to-expression modeling, cross-validation and choice of test and training samples. a) The goal is to predict expression of a gene G in conditions (i.e., samples) C_1_, C_2_, …, C_m_ using the expression of transcription factors TF_1_, TF_2_, …, TF_n_ in the same conditions. b) The schematic view of dividing conditions into training and test sets in cross-validation. A model is trained using the expression of TFs and gene G on the training set and is used to predict the gene expression in the test conditions. The predicted gene expression values (Ê) are compared to the real gene expression values (Ê) to provide a measure for the accuracy of the model. c,d) The illustration of different ways of dividing conditions into test and training sets. In this toy example, only two TFs are considered and each point represents a condition. In (c) test points are located close to training points, but in (d) test points are far from training points. Evaluating the performance of a model such as the one shown in (a) using the partitions shown in (c) or (d) may lead to different results.

The reasoning behind using CV to (indirectly) evaluate the accuracy of GRN reconstruction methods is that the relationships between TFs and their target genes remain mostly unchanged across conditions, and therefore a model correctly identifying TF-gene relationships based on the training set should make good predictions on the test set. We note that studies of context- or condition-specific regulatory relationships ^8,9^ do not adopt this evaluation strategy, and instead of learning a global GRN, a GRN per context is learned. In this study, our focus is only on methods for inference of global GRNs using transcriptomic data and CV-based methods for their evaluation. Cross-validation is a widely accepted approach for assessment of prediction (classification or regression) methods. Among the advantages of using CV is the fact that accuracy of two different models can be compared even if the models are of different complexity, e.g., if one model employs more TFs to explain the gene’s expression than another model. It is also an extremely valuable tool in bioinformatics, since experimental testing of predictions is often tedious and expensive and is typically done on a limited scale.

However, one problem with this popular approach is that CV results may depend on similarity of test and training sets. Consider first a contrived and extreme scenario where each data sample is present in many copies in the available dataset, so that in any random partition during CV (the standard way to construct training and test sets) a test sample is likely to have a copy of itself present in the training set. A supervised learning method may predict accurately on such a sample since it has already seen that sample during training. In this case, the high predictive accuracy of the model in the CV evaluation may simply be the result of this proximity between training and test points (e.g., by a successful adoption of the nearest-neighbor approach ^10^), and does not necessarily imply an accurate encapsulation of the input-output relationship in the trained model. (Note that this issue also occurs when using the leave-one-out strategy ^11^, another popular cross-validation method). In the context of GRN recovery, in which a data sample may be a single experimental condition represented by a vector of TF expression values, less extreme versions of the above scenario may manifest (Fig. 1c) through the presence of several biological replicates of the same conditions in the dataset. Thus, for the same reason as above, the accuracy estimated by CV on random partitions – as opposed to other partitioning approaches (Fig. 1d) – may be misleading and not generalize to more dissimilar experimental conditions. Put another way, while the goal of CV is to assess the ability of the method to predict the outcome on “unseen” samples, the line between “seen” and “unseen” samples may be blurred in standard CV on a typical gene expression dataset, as training and test sets may contain samples with extremely high similarity; this may in turn give us a false impression of predictive ability of the model and hence of the reliability of the inferred regulatory influences on the gene.

One might legitimately argue against the above critique: a supervised prediction method is only expected to learn how to predict on unseen samples that are drawn from the same distribution as training samples; an evaluation of its performance ought to respect this assumption, as is the case with CV on random partitions. We do not question this point of view in the general case, but only in contexts in which the model evaluated using CV is intended to elucidate feature-response relationships that are supposed to be generalizable beyond the characteristics of the dataset on which it was trained. For example, in global GRN reconstruction, training a prediction model is only a means to learn TF-gene regulatory relationships, which are represented by the parameters of the model; the actual prediction on unseen data is only indirectly relevant. If the CV is set up in a way resembling the above scenario, with test samples being similar to (e.g., biological replicates of) training samples, then a model may perform well on test data without accurately learning TF-gene regulatory relationships, yielding strong CV performance but not so strong GRN inference, as was observed in ^12^. Indeed, some previous studies have recognized this issue, in contexts related to GRNs and otherwise, requiring CV to be followed by ‘cross-condition’ assessments such as testing of models on data from unseen cell types ^13,14^.

In this study, we seek to explore the aforementioned problem and to identify and assess alternative strategies so as to avoid the pitfall mentioned above. More specifically, our goal in this study is not to propose a new GRN recovery algorithm, but to propose new CV strategies better capable of assessing the generalizability of such algorithms on unseen data. To this end, we first use clustering based cross-validation (CCV) – a non-standard approach to CV previously proposed in other contexts ^15–17^ – and illustrate that CCV provides a more realistic estimate of performance on unseen samples compared to random cross-validation (RCV), the standard method of CV. (Here, we specifically focus on the case of unseen samples being qualitatively distinct from training samples, in some objective sense, such as belonging to a distinct cell type.) In the CCV approach, we create CV partitions by first clustering the experimental conditions and including entire clusters of similar conditions as one CV fold. This strategy may be thought of as defining one group of trans-regulatory contexts as training data and testing a regression method trained on these contexts for its ability to predict gene expression in an entirely new regulatory context.

Noting that the main difference between CCV and RCV is that test samples in CCV tend to be distinct from training samples in the former, we next propose a score for the ‘distinctness’ of a test experimental condition (sample), based on a given set of training conditions. This score is independent of the model/algorithm that might be used for prediction. In addition, it is computed without the knowledge of the target gene’s expression levels in either training or test conditions, and is purely based on the predictor variables (TF expression values). We then generalize this to define a distinctness score for any given partition of the samples into training and test sets and show that gene expression prediction accuracy is highly (negatively) correlated with the distinctness score, confirming our expectation that it is easier to predict gene expression in a test condition that is similar to training conditions.

By comparing the performance of a LARS (Least Angle Regression) algorithm ^18^ on real transcriptomic data using RCV and CCV, we find that the RCV creates partitions for which the test folds are relatively easily predictable. On the other hand, relying on CCV for evaluation imposes several constraints. First, CCV inevitably inherits characteristics of the utilized clustering algorithm, such as dependency on the number of clusters, initialization, etc., and introduces the new problem of how to choose the clustering algorithm and its parameters, thereby adding subjectivity to the CV process. Second, a poor choice of clustering parameters may result in CCV not generating sufficiently distinct test/training partitions, thus failing to address the above-mentioned problem. To address these issues, we propose a CV approach based on simulated annealing (SACV) to construct any desired number of partitions spanning a spectrum of distinctness scores, allowing for a more controlled use of cross-validation for assessment and mutual comparisons of different methods of expression-to-expression modeling. Using this approach, we show that while the performance of two prediction algorithms (support vector regression (SVR) and Elastic Net) for expression modeling can be very similar for partitions with low distinctness score (i.e., with RCV), the difference between their performances becomes evident as the distinctness of partitions increases.

Our new CV strategy allows us to compare two algorithms not simply based on two numbers (i.e., their average RCV accuracy), but rather based on many such values obtained from a spectrum of partitions with different distinctness scores. This strategy makes the test/training distinctness an explicit and well-defined parameter of the evaluation process and enables us to evaluate the performance of different methods while considering the heterogeneity between the training set and the unseen dataset for which the method is intended. For instance, a method that performs better for small distinctness values may be more appropriate in cases where the unseen dataset is expected to be similar to the training set, while a method performing better for large distinctness values is appropriate when these two sets are expected to be dissimilar. Note that this feature of SACV has applications beyond the GRN recovery problem, as it can provide new insights on the performance of any classification/regression algorithm.

## RESULTS

### Overview of the gene expression prediction problem and cross-validation strategy

In a common formulation of the gene expression prediction problem, the goal is to predict a gene’s expression in a cellular condition, given the expression levels of all possible regulators (TFs) in that condition. We are given the expression levels ***E*** of (1) all TFs *T* and of (2) a gene of interest *g*, in a set of conditions *C*; from these data we must learn how to predict the expression of gene *g* from the expression of all TFs, in any condition, seen or unseen. (We assume here that *g* ∉ *T*.) We denote the expression of a TF *t* in condition *c* by *E*_*ct*_ and that of a gene *g* by *E*_*cg*_. The vector of all TF expression levels in condition *c*, i.e., {***E***_*ct*_}_*t*∈*T*_, is denoted by ***E***_*c*_. The gene expression prediction problem may thus be considered as a multivariable regression problem, where a model is trained to map the input vector ***E***_*c*_ to the scalar value ***E***_*cg*_.

The model must be able to accurately predict *E*_*cg*_ from ***E***_*c*_ even for *c* ∉ *C*, i.e., it must generalize well to predict accurately on unseen data; as such, model training must avoid over-fitting. Often, the number of conditions with available data (|C|) is in the tens or hundreds, and the number of TFs (|T|) is in the hundreds or thousands, i.e., the dimensionality of input vectors is of the same or greater order of magnitude as the number of training samples, making over-fitting a serious concern. This also means that the model evaluation has to be performed in a cross-validation setting, which may be loosely described as follows: assuming expression data are available for a set of conditions *C*, choose a subset of conditions *R* ⊆ *C*, called the ‘training conditions’, train the model using {(***E***_*c*_, ***E***_*cg*_)}_*c*∈*R*_, present the model with the TF expression vector ***E***_*c*_ for condition *c* ∈ *C* – *R*, called the ‘test condition’, and compare its prediction ***E***̂_*cg*_ to the known expression level ***E***_*cg*_ to assess the model’s accuracy (Fig. 1b). We call any given division of all conditions *C* into sets *R* and *C* – *R* (the training and test conditions) a ‘CV partition’. A ‘*K*-fold CV partition collection’ is defined as follows: all conditions in *C* are divided into *K* mutually exclusive and exhaustive subsets or ‘folds’ *S*_*1*_, *S*_*2*_, … *S*_*K*_ (often these folds are of equal or comparable sizes, though this is not an assumption in our treatment); each subset *S*_*i*_ is then used to define a CV partition by treating *S*_*i*_ as the set of test conditions and *R*_*i*_ = *C* – *S*_**i**_ as the training conditions. The collection of these *K* partitions is called a ‘*K*-fold CV partition collection’.

### Overview of gene expression data

In order to build and compare different expression-to-expression models and assess different evaluation strategies, we obtained mRNA expression data from The Cancer Genome Atlas (TCGA) corresponding to 12 cancers in the PanCan12 dataset ^19^ (see Methods). These served as the dataset for analyzing various cross-validation approaches. In addition, we obtained PanCan-normalized gene expression data from four other cancers from TCGA, to serve as ‘held-out’ data. Moreover, we obtained a list of human TFs from the Animal Transcription Factor Database (AnimalTFDB) ^20^, of which 1239 were present in this TCGA dataset. Since this dataset contains transcriptomic data from different biological contexts (i.e., cancer types), it allows us to explore the effect of different partitioning methods on the generalizability of the learned model to a new biological context.

In the PanCan12 dataset, one cancer type only contained 72 samples. In order to ensure that all cancer types are equally represented, we randomly selected 72 samples from all other cancer types, resulting in 864 samples in total. Before performing in-depth modeling of the dataset, we sought to visualize how expression varies across the different biological contexts. First, we represented each sample as a 1239-dimensional vector in which each entry corresponds to the expression of one TF. We used the multidimensional scaling (MDS) technique ^21^ to map these samples onto a two dimensional space (see Fig. 2). We noted that samples from different cancer types fall into distinct clusters based on the expression of their TFs, and most of the samples from each cancer type are placed in only one of these clusters. This simple visualization supports a key premise of our study – that transcriptomic samples, represented by their TF expression profiles, cluster by their biological context; as a result, assessing the performance of a model trained on data from one set of contexts using ‘distinct’ test samples (belonging to a different cluster) may be useful to evaluate its generalizability to unseen data from a biologically different context. Admittedly, not every cancer is a separate cluster in Fig. 2, reminding us that different biological contexts may not always have distinct and detectable signatures in the TF expression space.

**Figure 2.**
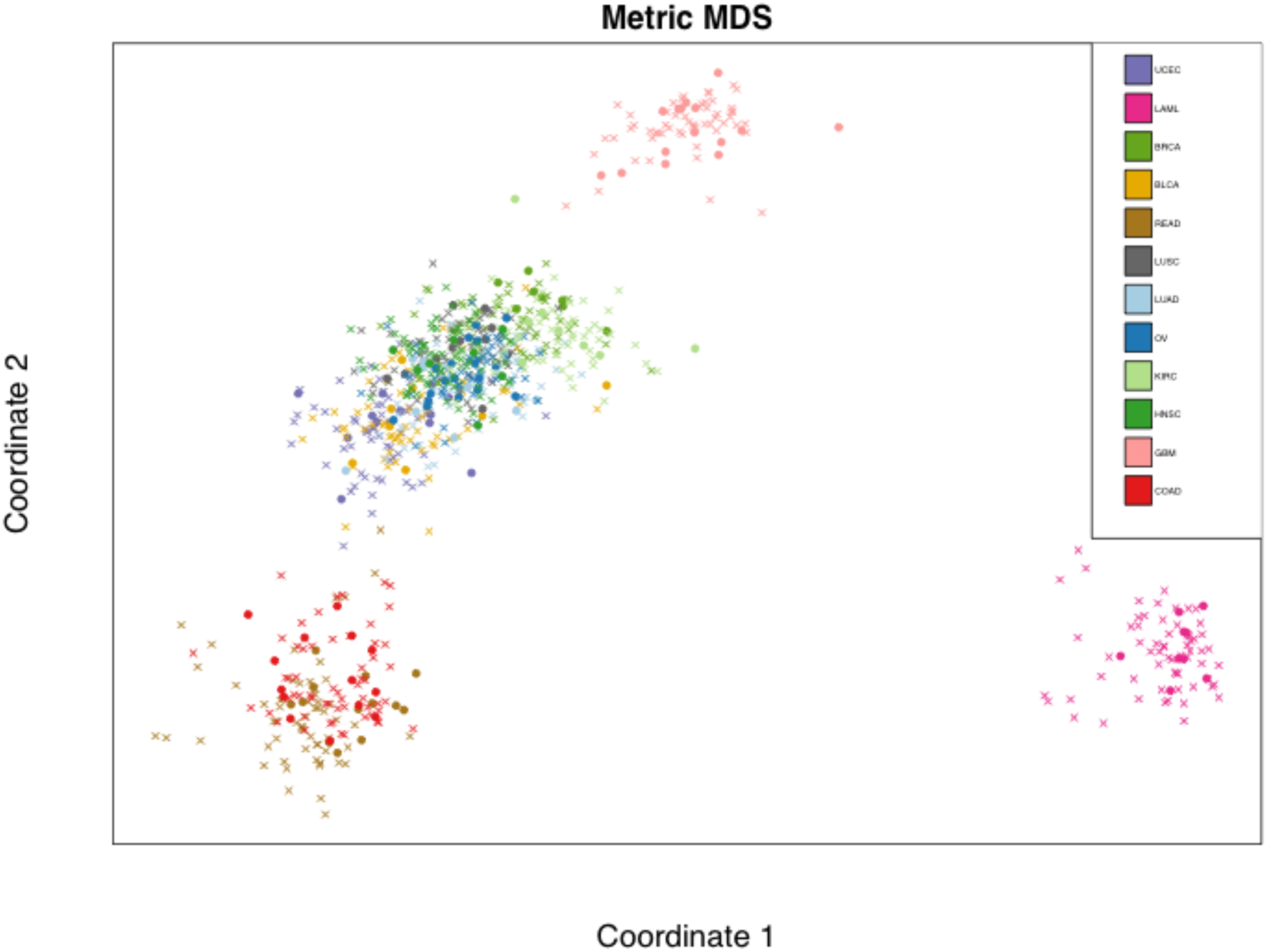
Samples from different cancer types and their random partition into training and test sets. Each sample (represented using a vector of 1239 TFs expressions) is mapped onto a 2dimensional space using multidimensional scaling. Samples of various cancer types of PanCan12 dataset are shown in different colors. Circles represent test samples (i.e., test conditions) and crosses represent training samples.

### Random and clustered cross-validation produce different estimates of error

Our goal was to model a gene’s expression in a sample based on the expression of 1239 TFs in that sample. To this end, we selected 500 target genes, which had the highest expression variance across the 864 PanCan samples. Following Chandrasekaran et al.^6^ we used the LARS regression method ^18^ as the gene expression prediction algorithm in our study. (LARS is a specialized linear regression algorithm suited for high dimensional data.)

To evaluate the prediction performance, we performed cross-validation using two approaches: RCV and CCV. In RCV, we randomly divided all conditions *C* into six equal folds, and defined each CV partition as a training set *R* of conditions in five of the six folds (720 samples) and a test set *S* = *C* – *R* comprising the remaining conditions. We repeated this procedure 10 times, obtaining 10 different 6-fold RCV partition collections. As was expected, RCV ignored the underlying distribution of the data and placed training and test samples close to each other. This is shown in Fig. 2 for one choice of training/test sets, and is reminiscent of the schematic view of Fig. 1C. In the CCV approach, we used k-means (k=6) clustering of the 864 samples based on their 1239-dimensional TF expression profiles to divide the samples into six subsets, which were then used to form training and test sets. This procedure was also repeated 10 times (with different initial seeds given to the k-means clustering algorithm), providing 10 different 6-fold CCV partition collections (see Methods for more details).

We evaluated the accuracy of LARS-based gene expression prediction for each of the 20 CV partition collections described above. Accuracy was measured using Pearson correlation (higher is better) between the predicted and the measured expression values of the 864 samples. Note that the predicted value of a sample is the value obtained using LARS when that sample is in the test set. Since each of the 500 genes represent a separate prediction task, we obtained a Pearson correlation coefficient (PCC) for each of these genes. Figure 3 shows the distribution of these 500 PCC values for each of the 20 partition collections. It is clear from these plots that the CCV provides a less optimistic evaluation of the LARS accuracy compared to RCV. This is also apparent in a direct comparison of accuracy assessed by the two cross-validation strategies for each gene separately (see Supplementary Fig. S1).

**Figure 3.**
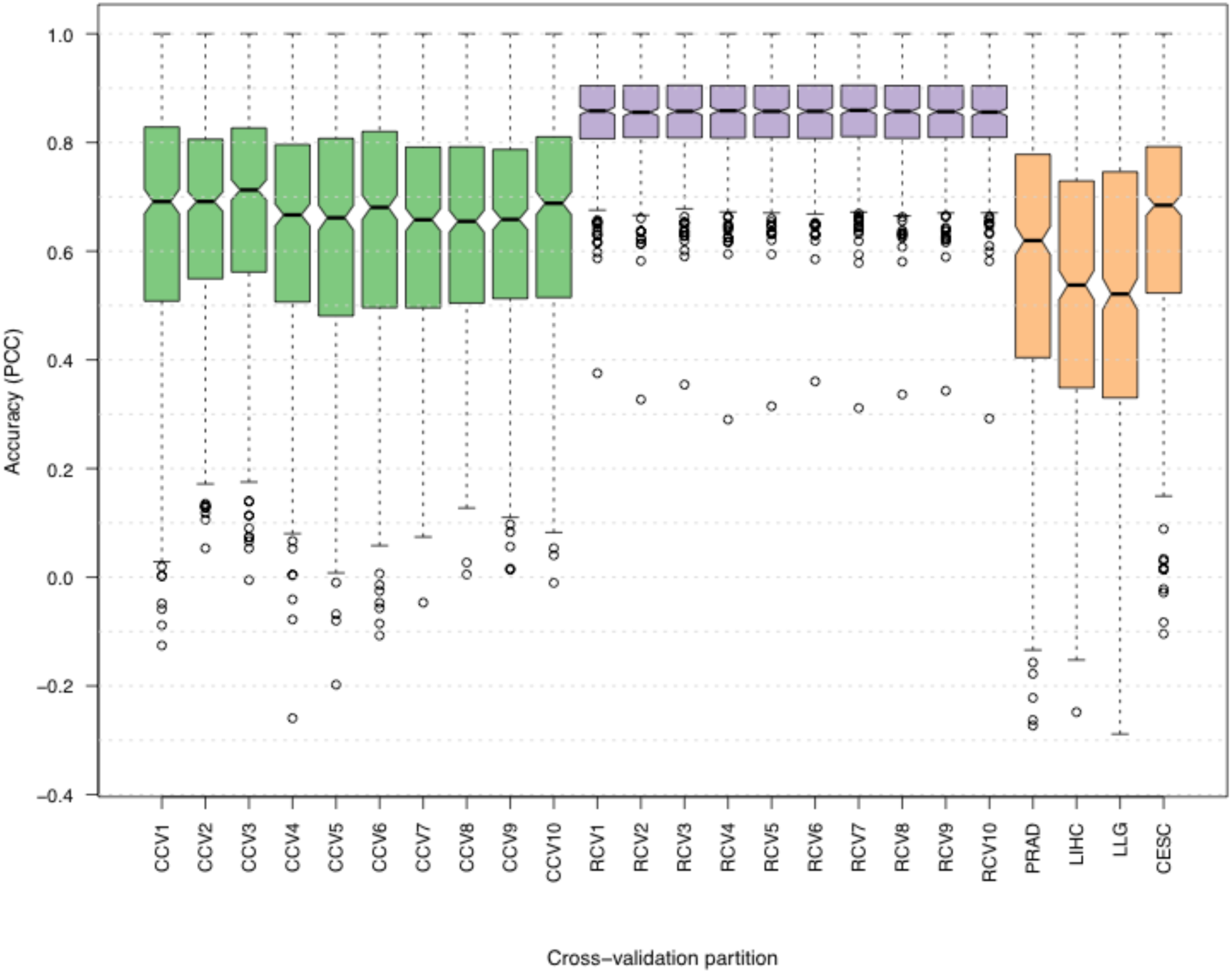
Cross-validation performance of the LARS regression model for gene expression prediction. Distribution of the Pearson correlation coefficients between predictions and real gene expression values over all test conditions. Each box plot shows the correlation calculated for 500 different genes. The ten boxplots labeled as CCV/RCV represent ten different clustering-based/Random 6-fold CV partition collections. Boxplots labeled as PRAD, LIHC, LLG, and CESC represent the performance of the regression method on conditions associated with cancer types not present in cross-validation studies. A higher Pearson correlation indicates higher accuracy.

Given these two contradictory assessments of the LARS prediction performance, we studied which one of the two CV strategies provides a more accurate estimate of the LARS prediction performance when a new set of samples corresponding to entirely different biological contexts is used as the test data. To this end, we formed four new test sets, each corresponding to one of the four different cancer types (PRAD, LIHC, LLG, CESC) that were not included in the PanCan12 dataset above. The 864 samples from the PanCan12 were used as the training set. Figure 3 shows that RCV produces an over-optimistic estimate of the accuracy, while CCV provides a more realistic estimate of the accuracy on these four test sets that are qualitatively different (i.e., different cancer types) from the training data.

### A new measure to quantify the distinctness of training and test sets

Although CCV provides a more realistic estimation of accuracy compared to RCV on datasets dissimilar to the training set, it relies on a clustering algorithm and is limited by the latter’s properties. Choice of the number of clusters, the clustering criteria driving the algorithm, or even its initialization may severely affect the algorithm’s output, and hence the performance estimate obtained using CCV. As a result, CCV introduces the new problem of how to choose the clustering algorithm and its parameters. Moreover, the clustering algorithm used in CCV may not generate sufficiently distinct test/training sets. When it does produce distinct sets, the clusters may be at different distances from each other and averaging the estimated performance over all clusters may not be a reliable estimate of performance on new samples.

To address the above issues, we first sought to quantify how distinct training and test sets are from each other. Doing so should allow us to report cross-validation accuracy under controlled levels of training/test set distinctness, e.g., by selecting as test sets those clusters that are distinct from the remaining clusters (used as training sets), and discarding those partitions that lead to relatively similar training and test sets. Explicitly quantifying the distinctness of training and test sets may also allow us to directly construct test/training sets of a specified distinctness, bypassing the clustering step altogether, as discussed later in the manuscript.

We first assigned to each test sample *i* (represented by its TF expression vector ***E***_*i*_) a score *η* capturing its distinctness from the training set *R*:

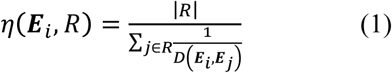

where *D*(***E***_*i*_, ***E***_*j*_) is the min-max normalized Euclidian distance (see Methods) between TF expression vectors ***E***_*i*_ and ***E***_*j*_, representing the test condition ***i*** and training condition ***j***, respectively. Our choice of the harmonic mean in equation (1) reflects our belief that the presence of even one or few training sample(s) very similar to the test sample (i.e., relatively close TF expression vectors) is sufficient to violate the distinctness: the harmonic mean is dominated by the distance of the closest training samples to the test sample. We note that the distinctness of a test sample is estimated without the knowledge of the gene expression level *E*_*g*_ of the target gene.

Given this definition, we evaluated the distinctness of test samples from training sets for the 10 RCV and 10 CCV partition collections described above. Figure 4a shows the distribution of the distinctness scores of the test samples (scored against corresponding training sets) for each of these 6-fold partition collections. It is clear from this figure that the distinctness scores in CCV partitions are generally greater than in the RCV partitions, which is expected as the k-means clustering algorithm used by the CCV procedure identifies groups of conditions that are close to one another and distant from the rest.

**Figure 4.**
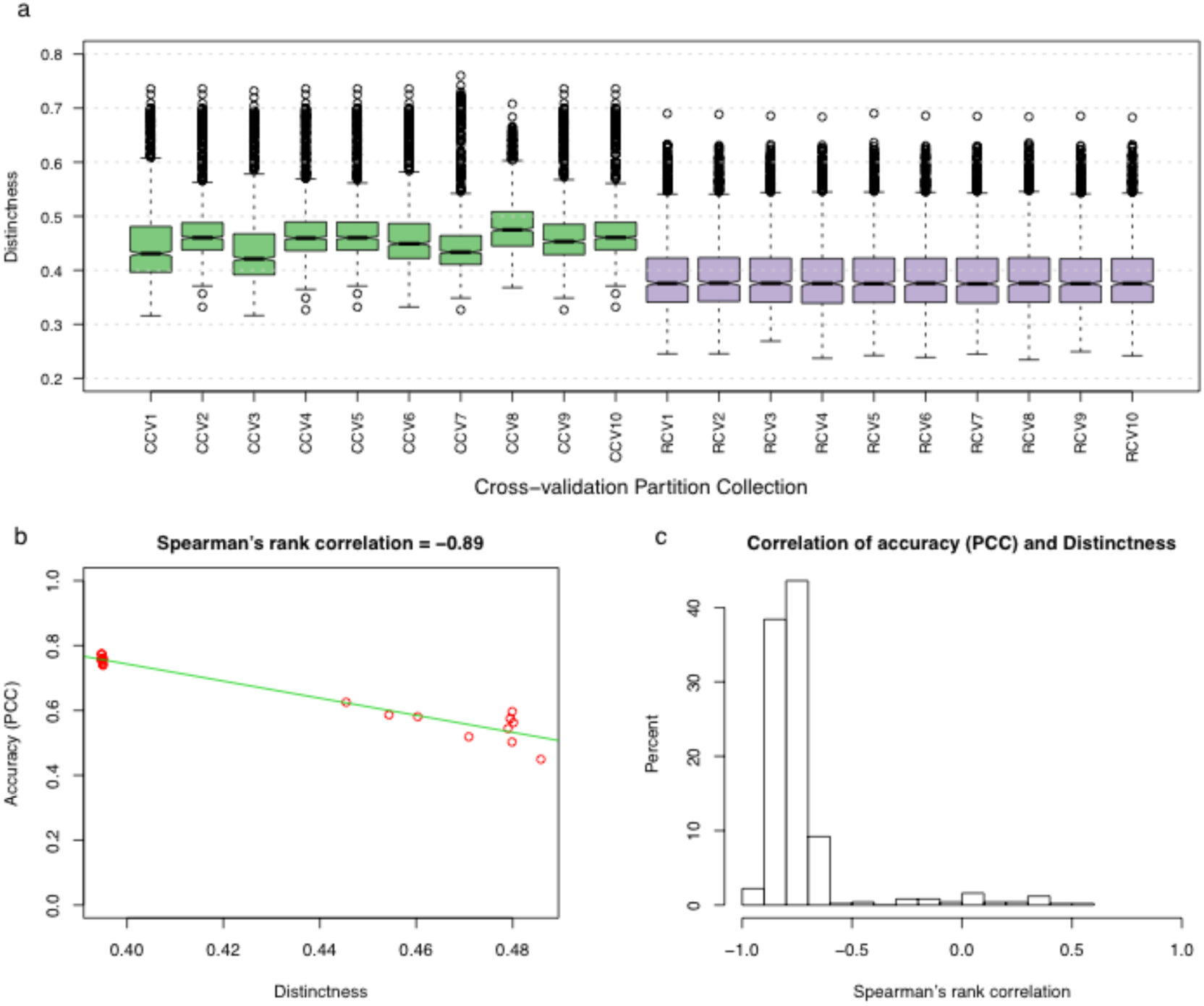
Distinctness score of test conditions in clustered vs. random cross-validation schemes. a) The distribution of distinctness score under various cross-validation schemes. Each boxplot represents distinctness scores of 864 conditions corresponding to a CV partition collection. b) The relationship between distinctness and performance of expression prediction for one gene. The plot shows the PCC values against the values of the distinctness scores for all 20 CV partition collections. The Spearman’s rank correlation between distinctness scores and PCC scores is equal to −0.89 for this gene. c) Histogram of the Spearman’s rank correlation values between the distinctness and the prediction accuracy for 500 genes. Each correlation value is calculated as shown in Figure 4b.

Based on the above definition, we next defined the distinctness score of an entire *K*-fold CV partition collection *P*, described by the test sets 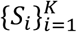 and associated training sets 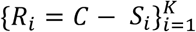, as the average distinctness of all test samples across all partitions:

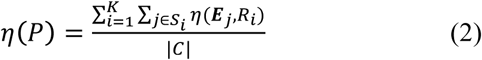

Note that this distinctness measure also only depends on the samples’ TF expression and does not depend on the target gene’s expression.

### Gene expression prediction accuracy estimated by cross-validation is highly correlated with distinctness score of test conditions

Generalizing from our observation that CCV produces less optimistic performance estimates than RCV, due to test sets that are by construction distinct from training conditions, we hypothesized that an appropriately defined distinctness score of a (test, training) partition is negatively correlated with the performance of the trained regression model. To test whether the distinctness score we defined above satisfies this condition, we evaluated the distinctness of the 10 RCV and the 10 CCV partition collections (described above), and compared the distinctness values to the PCC measures of accuracy. Figure 4b shows the relationship between distinctness of a CV partition collection and accuracy of LARS regression method on that partition collection for one gene as an example. It is clear that CV accuracy (for the task of predicting this gene’s expression) decreases with increasing distinctness scores, with a Spearman’s rank correlation of −0.89. Figure 4c summarizes this information for all 500 genes, using the histogram of correlation coefficients between the distinctness score values and the estimated performance values of the LARS predictions. These results supported our expectation of a negative correlation between distinctness score and prediction accuracy: distinctness of a CV partition collection, as estimated by the score *η*(*P*), has a correlation of −0.7 or less with prediction accuracy for over 84% of the 500 genes.

### Construction of cross-validation partitions with varying distinctness scores

As we previously demonstrated (Fig. 4a), test and training sets obtained using CCV are more distinct than the ones obtained using RCV. However, there is a limited number of ways to generate such partitions since they rely on the results of a clustering algorithm. We thus asked whether it was possible to obtain a set of CV partitions with varying degrees of distinctness, without relying on a clustering method. To this end, we implemented a simulated annealing algorithm ^22^ to sample partitions of gradually increasing distinctness score (see Methods for details). In each step, this algorithm generates a new CV partition with a higher distinctness score. Figure 5a shows the distribution of the distinctness scores of the test samples in the CV partitions generated using this algorithm in 30 steps, each containing 200 test samples. The CV partitions represented in this figure are uniformly sampled from all of the CV partitions generated by the simulated annealing algorithm (see methods). The median distinctness score of each CV partition, across the 200 test samples, ranges between ~0.4 (which matches the average distinctness of the RCV partition collections in Fig. 4a) and ~0.55 (which is greater than the average distinctness of the CCV partition collections in Fig. 4a). This shows that our procedure can be used to generate a fixed size test set with user-specified distinctness score in the range spanning from that of RCV partitions at the low end and CCV partitions at the high end, and even higher.

**Figure 5.**
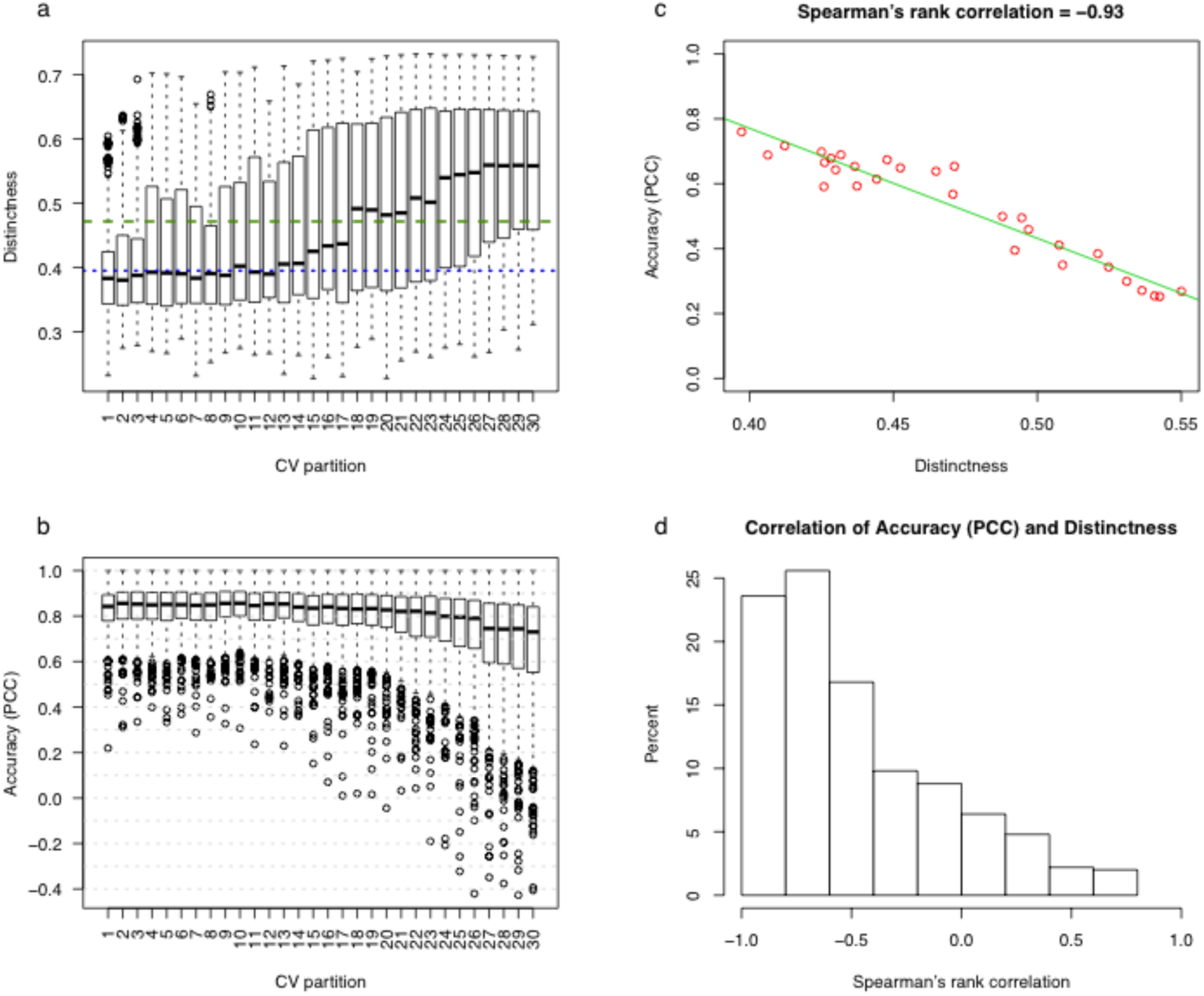
Systematic generation of cross-validation partitions with gradually increasing distinctness score using simulated annealing. a) The distribution of distinctness scores of 200 conditions in each CV partition generated using simulated annealing. The dashed and dotted lines show the average distinctness score of the 10 CCV and the 10 RCV partition collections, respectively. b) The distribution of the test set prediction accuracies for 500 genes. The prediction accuracy is measured using the PCC of model-predicted expression values and real expression values. c) The accuracy of predicting test set expression of one gene for the 30 cv partitions versus the distinctness score of the CV partitions. d) The histogram of Spearman’s rank correlation coefficient between the prediction accuracy and average distinctness value, across the 30 CV partitions. The histogram includes one correlation value for each of the 500 genes. Each correlation value is calculated as shown in Figure 5c.

Figure 5b shows that the cross-validation accuracy (measured using PCC) of LARS decreases as successive steps of the simulated annealing algorithm generate CV partitions of increasing distinctness scores. This behavior is in agreement with the trend observed in Fig. 4 but at a finer granularity. Figure 5c shows the correlation between LARS accuracy and distinctness score of CV partitions generated by the algorithm, for a randomly selected gene. Finally, Fig. 5d confirms this observation more globally – the Spearman’s rank correlation between distinctness scores and LARS performance, across the 30 CV partitions of Fig. 5a, is below −0.4 for 66% of the 500 genes. Supplementary Figure S2 shows this relationship for several randomly selected genes.

### More distinct cross-validation partitions can better reveal performance differences between prediction methods

We noted above (Fig. 5) that CV partitions with larger distinctness scores result in poorer performance estimates for the LARS prediction method. We sought to determine whether this trend is more conspicuous for one prediction method than for another. If so, then CV partitions constructed with relatively high distinctness scores may be better suited to discriminate between two (or more) prediction methods, and lead to different conclusions from comparative evaluation of those methods than CV based on random partitions.

To test for the above possibility, we used the previously constructed simulated annealing based CV partitions, which comprised of 30 partitions each containing 200 test samples and 684 training samples. (Thus, 30 CV partitions of varying distinctness scores were used.) We evaluated three different expression prediction methods for their ability to predict the expression of the 500 selected genes, using these 30 CV partitions (Supplementary Figs. S3 and S4). These methods include LARS, Elastic Net, and support vector regression (SVR). Figure 6 shows the performance differences (measured in root-mean-square deviation, RMSD) between SVR and Elastic Net, as a distribution over the 500 genes, for each CV partitions. These two methods have a similar performance on partitions with smaller distinctness score, with SVR performing slightly better (smaller RMSD). However, in the last six partitions, which have the highest distinctness scores, Elastic Net clearly outperforms SVR. These results show that the comparison between two gene expression prediction methods can be more informative if we use CV partitions spanning a spectrum of distinctness scores, resembling unseen data of various degrees of similarity to the training set, which can be produced by our proposed simulated annealing algorithm.

**Figure 6.**
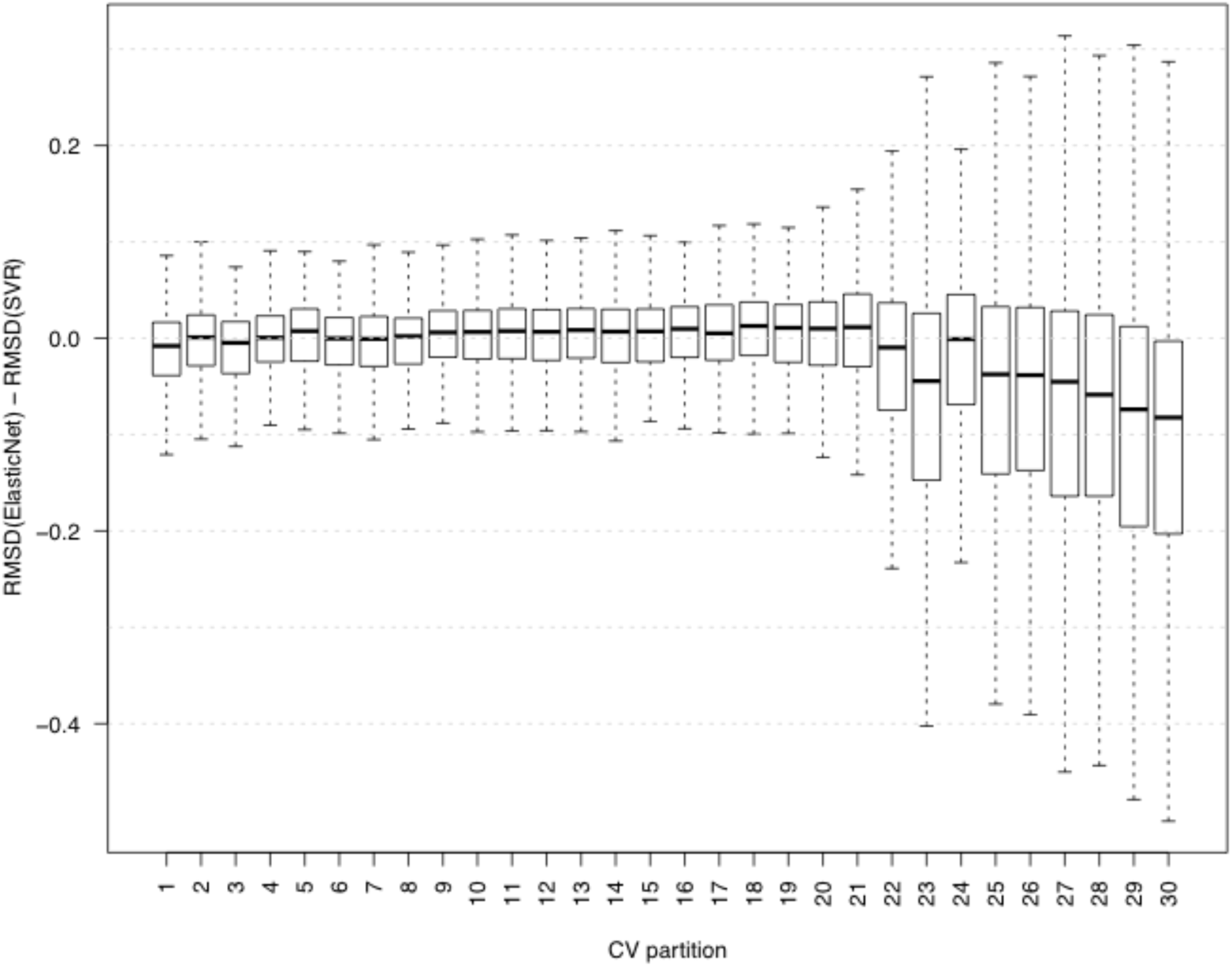
Comparing the performance of SVR and Elastic Net on CV partitions with different distinctness scores. Each boxplot shows the distribution of prediction accuracy differences between SVR and Elastic Net on 500 genes. The CV partitions are generated using simulated annealing. Outliers are not shown in this plot.

## DISCUSSION

Cross-validation is an extremely valuable tool for comparing the performance of diverse methods, especially in case of models with different number of parameters. However, such CV-based evaluation strategies must be properly designed for a particular application in order to result in correct insights and conclusions. For example, one of the important applications of gene expression prediction is to predict gene expression in a new and distinct regulatory context (e.g., different tissue or disease type). In such cases, one needs to evaluate the performance generalizability of gene expression prediction methods to new samples that may not be well represented by the training samples. As we showed in this study, random cross-validation provides an over-optimistic performance evaluation, which does not generalize to new biological contexts. This is due to the fact that RCV may place samples with high similarity in both the training and test sets, blurring the line between seen and unseen samples. This may happen to different extents in different datasets, which makes it difficult to interpret, in any absolute sense, the performance estimated by CV on different datasets: two studies may report the same CV prediction accuracies, but depending on the heterogeneity of their datasets the reported accuracy in one study may be more generalizable than the other.

To address these issues, we first proposed a clustering-based CV, which allows us to obtain a more realistic estimation of error on distinct test samples. On the other hand, since CCV relies on a clustering algorithm, the choice of the algorithm and its parameters may play an important role on the obtained clusters and hence the performance evaluation. In addition, the similarity of the unseen data to the training set is a significant factor in determining which one of RCV or CCV strategies generates more accurate prediction estimates: if the unseen data is well represented in the training set, RCV may be more accurate. To address these issues, we introduced a simulated annealing approach, which is not limited by the characteristics of any particular clustering algorithm and allows us to construct test/training partitions with desired degrees of distinctness (a measure that quantifies how similar a test set and a training set are to each other). In the context of CV, the distinctness score can be thought of as an extra parameter which allows us to compare two prediction algorithms more accurately: instead of obtaining average accuracy performance of each algorithm and comparing these two numbers, one can compare their performance on test sets spanning a spectrum of distinctness scores, resembling unseen data of various degrees of similarity to the training set. As we showed in this study, two prediction methods (SVR and Elastic Net) had similar performance on less distinct partitions and a RCV evaluation would have concluded that these methods approximately perform equally well. On the other hand, more distinct partitions revealed that such a conclusion does not hold if the test sets are more distinct from the training sets and Elastic Net performs better.

It is worth pointing out that the ability of a training dataset to provide accurate estimates of the prediction performance on unseen data depends on the heterogeneity of the training data as well: in a homogeneous dataset comprising of mostly similar conditions, any CV technique (even SACV) can produce partitions with only a limited range of distinctness scores. As a result, such a dataset may be most suited to generate context-specific predictions. However, even in such a case, utilizing SACV to generate partitions with a spectrum of distinctness scores may provide a better means to compare prediction performance of different algorithms.

Another important point is that while the distinctness score is negatively correlated with the performance, the actual value of performance measure also depends on the size of the test sets. As a result, when comparing two prediction methods, one must ensure that the test set sizes are equal. This is yet another benefit of using SACV in comparing methods (specially from different studies), since it can generate partitions with varying distinctness for any desired test set size, allowing to compare performance accuracies while fixing the test size and the distinctness score.

## METHODS

### Data Collection

We obtained mRNA expression data from The Cancer Genome Atlas (TCGA) corresponding to 12 cancers in the PanCan12 dataset ^19^: glioblastoma multiformae (GBM), lymphoblastic acute myeloid leukemia (LAML), head and neck squamous carcinoma (HNSC), lung adenocarcinoma (LUAD), lung squamous carcinoma (LUSC), breast carcinoma (BRCA), kidney renal clear-cell carcinoma (KIRC), ovarian carcinoma (OV), bladder carcinoma (BLCA), colon adenocarcinoma (COAD), uterine cervical and endometrial carcinoma (UCEC) and rectal adenocarcinoma (READ). This dataset includes RNA-seq expression profiles of primary tumor samples in 3599 individuals and was downloaded from ‘https://xenabrowser.net/’. In addition, we obtained PanCan-normalized gene expression of four other cancers from TCGA: prostate adenocarcinoma (PRAD, 550 samples), liver hepatocellular carcinoma (LIHC, 423 samples), brain lower grade glioma (LLG, 530 samples), and cervical squamous cell carcinoma and endocervical adenocarcinoma (CESC, 308 samples). Moreover, we obtained a list of human TFs from the Animal Transcription Factor Database (AnimalTFDB) ^20^, of which 1239 were present in this TCGA dataset.

### Clustered cross-validation

We used k-means clustering (k=6) in order to construct clustered CV partitions. Clustering was performed on 864 samples included in our study. Each output cluster was then used as one CV fold. This procedure was repeated 10 times with different initial seeds for the k-means algorithm, resulting in 10 different CCV partition collections.

### Min-max normalized distance

To calculate the min-max normalized Euclidean distance between two conditions (TF vectors), we first calculated the Euclidean distances between all pairs of conditions. Let *D*_*min*_ and *D*_*max*_ denote the minimum and maximum of these Euclidean distances. Then, given the Euclidean distance of two conditions *D*, their min-max normalized Euclidean distance is calculated as

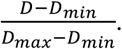

### Regression algorithms

LARS, Elastic Net, and support vector regression (SVR) with Gaussian kernel were performed in MATLAB^™^ using the expression of TFs in each condition as predictors of gene expression in that condition. The hyperparameters of LARS and Elastic Net were chosen using an inner 5-fold CV in the training set. Gene expression data used as the input to the regression algorithm was z-score normalized.

### Simulated Annealing (SA)

SA ^22^ was performed to generate CV partitions with varying degrees of distinctness; hence we used the defined distinctness score of a CV partition as the objective function. Starting with a random choice of conditions as the test fold and initial temperature equal to 1, in each iteration at a specific temperature, one condition from the test set was substituted with a condition in the training set. The distinctness of the new CV partition was evaluated and the substitution was accepted with a probability calculated based on the change in distinctness of the CV partition and the SA temperature of that step. After 500 iterations, the temperature was decreased by a factor of 0.98. This substitution procedure was repeated until the temperature reached the value of 1e-14, resulting in the 73423 accepted CV partitions. All of the accepted CV partitions from all iterations (at all different temperatures) were recorded. The first 50000 CV partitions produced by the SA algorithm were discarded and a total of 30 CV partitions with varying degrees of distinctness were uniformly sampled from the remaining SA-produced CV partitions.

## Data Availability

The datasets analyzed during the current study are available in the UCSC Xena repository, [https://tcga.xenahubs.net/download/TCGA.PANCAN12.sampleMap/PanCan12.3602-corrected-v3_syn1715755.gz1].

## Author Contribution

SS, SDZ conceived and designed the study. STB, AE designed and performed analyses. All authors wrote and reviewed the manuscript. STB prepared figures.

